# Evolutionary branching in multi-level selection models

**DOI:** 10.1101/2024.06.21.600152

**Authors:** Burton Simon, Yaroslav Ispolatov, Michael Doebeli

## Abstract

We study a model of group-structured populations featuring individual-level birth and death events, and group-level fission and extinction events. Individuals play games within their groups, while groups play games against other groups. Pay-offs from individual-level games affect birth rates of individuals, and payoffs from group-level games affect group extinction rates. We focus on the the evolutionary dynamics of continuous traits with particular emphasis on the phenomenon of evolutionary diversification. Specifically, we consider two-level processes in which individuals and groups play continuous snowdrift or prisoner’s dilemma games. Individual game strategies evolve due to selection pressure from both the individual and group level interactions. The resulting evolutionary dynamics turns out to be very complex, including branching and type-diversification at one level or the other. We observe that a weaker selection pressure at the individual level results in more adaptable groups and sometimes group-level branching. Stronger individual-level selection leads to more effective adaptation within each group while preventing the groups from optimizing their strategies for group-level games.

## 1 Introduction

There are numerous examples where individuals living in groups evolve behavioral and phenotypic patterns distinct from their solitary peers. Group selection mechanisms are often invoked to explain the evolution of cooperation in its most general form (e.g. [Traulsen and Nowak, 2006]) and in more specific examples of evolution of distinct individual and group strategies due to competitive pressure [Jackson, 1977], predation and evasion of being preyed on [Lang and Farine, 2017, Hebblewhite and Pletscher, 2002], and even the evolution of group-forming itself, e.g. in the form of monogamy vs polygamy in primates and humans [French et al., 2018].

Evidently, the different and often contrasting individual and group strategies evolve due to different selection pressures at the two levels in those scenarios. Such differences often emerge despite the underlying similarity of the fundamental evolutionary trend to increase birth rates and decrease death rates. Furthermore, the broadly understood external conditions are often the same for the evolution of distinct traits in solitary and in-group individuals. The degree of individual involvement and thus the relative evolutionary importance of the group activity varies between species, such as weak and often temporal associations in a deer herd vs. permanent distribution of roles in a colony of honeybees [Seeley, 1997]. So it is interesting to examine how the solitary and group strategies evolve depending on the degree of importance of events at each level. The relative importance of individual-level and group-level events should determine which strategies eventually dominate [Aoki, 1982, Kimura, 1983].

Another interesting question is whether adaptation at the individual level is beneficial or detrimental to adaptation at the group-level, and vice versa [Okasha and Paternotte, 2012, Shelton and Michod, 2020]. For example, for the standard problem of the evolution of cooperation by group selection [Traulsen and Nowak, 2006], adaptation at the individual level is by assumption detrimental for groups. Intuitively, a stronger selective pressure at either the individual or the group level should result in faster and more pronounced evolutionary adaptation at the corresponding level, yet it is unclear in general how this plays out in more complicated evolutionary scenarios, such as the ones considered in this paper.

Questions like these about group-structured populations can be addressed via mathematical models that take state-dependent birth and death rates at both levels explicitly into account. For example, the partial differential equation models of group-structured populations developed in [Luo, 2014, Cooney, 2019, Cooney, 2020, Cooney and Mori, 2022] have been used to study a number of interesting questions about group-structured populations, including the evolution of cooperation, as well as transitions of the dominant selection regime from the individual to the group level or vice versa. However, these models do not address what happens when there is a continuum of individual-level and group-level phenotypes. We will show here that continuous phenotypes in group-structured populations can lead to complicated evolutionary outcomes that have not been reported before in the literature. Furthermore, an assumption common to partial differential equation models is that groups reproduce by “cloning” themselves, an event that does not increase the variability of group-level phenotypes, and is not always biologically realistic. Here we assume groups reproduce via fissioning events, which do increase the variability of group-level phenotypes, and which may be more realistic.

In this paper we attempt to shed some light on these subjects using examples of continuously varying strategies in evolutionary games. We examine what happens when the individual-level and group-level selection pressures favour different trait distributions. Our main focus is on phenomenon of evolutionary diversification, or evolutionary branching [Geritz et al., 1997] [Dieckmann and Doebeli, 1999]. Generally speaking, during evolutionary branching an initially monomorphic population splits into two distinct and diverging phenotypic clusters due to frequency-dependent selection. This phenomenon is known to occur in continuous evolutionary games, such as the continuous snowdrift game [Doebeli et al., 2004]. At the individual level, branching implies the evolutionary emergence of two distinct strategic clusters that coexist at evolutionary equilibrium: one consisting of low-valued strategies (defectors), and one consisting of high-valued strategies (cooperators). At the group level, evolutionary branching of continuous traits has rarely been described. If it does happen, branching would imply the evolutionary emergence of two distinct types of groups, one consisting mainly of defectors, and the other consisting mainly of cooperators. In particular, in this case each group in the population would be approximately monomorphic, and hence the within-group population would not undergo branching. Therefore, individual-level branching and group-level branching are essentially mutually exclusive, and the level at which branching will occur will depend on the relative strengths of individual and group selection.

Here we first show that evolutionary branching at the group level is indeed possible when the groups play a continuous snowdrift game. Since branching at one level interferes with branching at the other, the evolutionary dynamics are complicated when the snowdrift game is played at both levels, and we investigate how the relative strength of selection at the individual and group levels affect the evolutionary dynamics.

Besides using the snowdrift game at both levels, we also investigate the classical conflict of defection at the individual level and cooperation at the group level by assuming an individual-level game that favors defectors (con-tinuous prisoner’s dilemma game) but a group-level game that favors evolutionary branching (continuous snowdrift game). This is somewhat similar to the usual setup when studying the evolution of cooperation by group selection, except that traditionally, no game structure is present at the group level (instead it is simply assumed that more cooperative groups “do better”). The relative strengths of the games at the two levels determines whether the evolutionary forces at the individual-level are dominant (the equilibrium is mostly defectors), or if the evolutionary forces at the group level are dominant (with two distinct types of groups at equilibrium), or if the equilibrium is some sort of compromise.

Our model of group structured populations will be described mathematically in the next section. It is similar to the models used to study group selection in [Simon, 2010, Simon et al., 2013, Simon and Pilosov, 2016]:

- Individuals play games against their group mates; and groups play games against other groups.
- Individual birth rates depend on payoff in the within-group game (higher payoffs lead to higher birth rates). Individual death rates depend on the size of the group.
- Group reproduction is by (random) fission. The fission rate depends on the group size. Groups die by extinction. Extinction rates depend on payoffs in the group-level game (higher payoffs lead to lower extinction rates).

To study individual-level vs group level branching, we use the same basic continuous snowdrift game at both levels, but modify the individual-level game by multiplying the payoff function by a scaling factor, *s*, so that we can vary the relative strengths of the games at the two levels. We then use numerical simulations to investigate the effects of varying *s* on the evolutionary dynamics. A similar approach is used in the last example, in which individual selection favours defection, but group selection favours evolutionary branching. Overall, our study is among the first to consider two-level evolutionary processes with continuous games at the group level, and to show that evolutionary branching can occur due to group selection.

## 2 The Model

We study a group structured population where each individual has a type *x* ∈ [0, 1] which does not change during its lifetime. An individual’s type determines the strategy it plays in a within-group game. We will consider games here, such as continuous snowdrift game and continuous prisoner’s dilemma game, where *x* measures the level of cooperation, with *x* = 1 being the highest level of cooperation and *x* = 0 being the lowest. The groups also play games against each other, not necessarily the same game the individuals play, but also a game involving levels of cooperation. The strategy a group uses in the group-level game is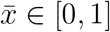, the average of the individual types in the group.

Although an individual’s level of cooperation does not change during its lifetime, a group’s level of cooperation can change during its lifetime due to births and deaths of individuals in the group. (Groups are born in fission events, and disappear in fission or extinction events, so a group’s lifetime is the span between those events.) When an individual gives birth, the offspring’s type is either the same as that of the parent individual (with probability 1 − *q*), or differs by a small Normally distributed amount (a mutation occurring with probability *q*). A mutant with a type outside the interval [0, 1] dies.

An individual’s birth rate depends on its expected payoff in the game with all other individuals in its group, which may change during its lifetime due to changes in the composition of its group. An individual’s death rate is proportional to the size of its group, so a group cannot grow without bound even without any group-level events.

Groups change in time due to internal births and deaths (individual-level events). They can also change suddenly due to group-level events. Groups can fission, where they break randomly into two pieces, and groups can die of extinction. A group’s fission rate is proportional to its size. A group’s extinction rate depends on the average payoff it earns in the between-group game and also on the number of groups in the population (which assures the number of groups cannot grow without bound).

The model just described can be formalized as a continuous-time Markov chain, which will be described next. However, the Markov chain we describe is analytically intractable due to its complicated state space, and even an exact simulation runs slowly when the groups are big, due to the very large numbers of individual-level birth and death events. Our numerical technique for analyzing the model approximates the within-group population dynamics by a deterministic process (a PDE), while retaining the stochastic fission and extinction processes at the group level. This hybrid approach, Simon and Nielsen (2012), will be explained and justified below.

### 2.1 Specifying the 2-level Markov chain

In order to specify the Markovian model exactly, we need the state-dependent rate functions for all the events that can change the state of the population. In the model there are four kinds of events: two individual-level events, births and deaths; and two group-level events, fission and extinction. Mutation, when it occurs, is part of the birth event.

A group is specified by the set of individuals it contains. Thus, the state of a group has the form *ξ* = {*x*_1_, *x*_2_, …, *x*_*n*_}, where *n* is the number of individuals in the group, and *x*_*i*_ ∈ [0, 1] is the type of the *i*th individual. The state of a group changes over time (including the variable *n*) due to births (with or without mutations) and deaths. The state of the two-level process has the form Ξ = {*ξ*_1_, *ξ*_2_, …, *ξ*_*N*_}, where *N* is the number of groups and *ξ*_*j*_ is the state of the *j*th group.

The continuous time Markov chain Ξ(*t*), *t* ≥ 0 is difficult to study analytically because (among other things) the number of groups in the population and the identities of the groups change in time due to fissions (group births) and extinctions (group deaths); while at the same time, each group changes in size and composition due to births and deaths of individuals. The state space for Ξ(*t*) therefore does not have a simple structure. We will address this difficulty after the Markov chain is properly defined.

Individuals play a game with their group mates with payoff to *x* in a game against *y* defined as *P*_*I*_(*x, y*). The group-level game has 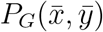 payoff to a group with average cooperation level 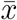 against a group with average cooperation level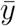.

The birth rate for the *i*th individual in a *ξ*-group (where *ξ* = (*x*_1_, …, *x*_*n*_)) has the form,

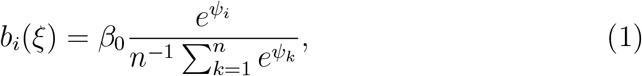

where

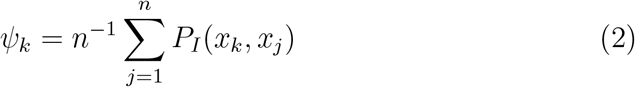

is the average game payoff to individual *k* from a game against a random opponent.

Two comments are in order regarding this choice for how the game affects birth rates at the individual level. First, since in general game payoffs can be positive or negative, the game payoffs are exponentiated so that only positive numbers enter the calculation. Second, the expression for the birth rates ensures that the total birth rate in a within-group population, i.e., the sum over all individual birth rates, is equal to *β*_0_*n*, and hence is independent of the game. Thus, the population dynamics of the total within-group population is decoupled from the game, which facilitates choosing appropriate population dynamics regardless of the choice of game. Without loss of generality, in the following the constant, *β*_0_, is fixed at *β*_0_ = 1 in all our examples.

Every individual in a *ξ*-group has the same death rate

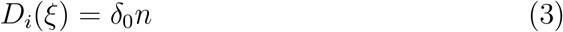

proportional to the group size. The death rate parameter, *δ*_0_, can be used as a simple way to control the average number of individuals per group in the model.

The fission rate of a *ξ*-group is

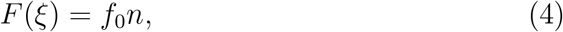

i.e., proportional to its size. When a group fissions, it breaks into two pieces, with every outcome equally likely (a uniform fission density [Simon, 2010]).

The extinction rate of the *i*th group in population Ξ = {*ξ*_1_, *ξ*_2_, …, *ξ*_*N*_} is

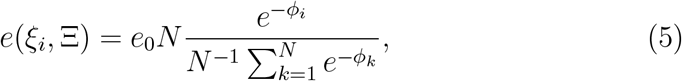

where

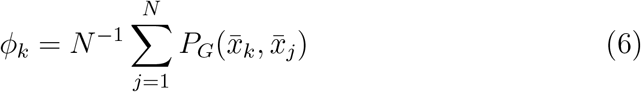

is the *i*th group’s expected payoff in the group-level game against all groups. As in the case of birth rates at the individual level, the average group game payoffs are exponentiated, but this time the exponents are negative since higher payoffs yield lower extinction rates. The group extinction rate function (5) plays a crucial role in our model. Since the fission rate (4) only depends on the group size, it is extinctions that must keep the number of groups in check, and impart advantages or disadvantages to groups that do well or poorly in the group-level game. The extinction rate is proportional to the number of groups in the population, *N*, so the population of groups cannot grow without bound. It is analogous to the individual death rate function in that regard.

As for birth rates at the individual level, the expression for the group extinction rates ensures that the total extinction rate in the population of groups, i.e., the sum over all group extinction rates, is equal to *e*_0_*N* ^2^, and hence is independent of the game played by the groups. Thus, the population dynamics of the total population of groups is decoupled from the group game, which again facilitates choosing appropriate population dynamics regardless of the choice of game.

As we mentioned above, the continuous-time Markov chain we have defined is analytically intractable. In theory it could be simulated exactly using e.g. the Gillespie algorithm, but when groups are large, the exact simulation becomes very slow due to the large number of births and deaths that can occur. We therefore chose to use the “hybrid” method for simulating group-structured populations described in [Simon and Nielsen, 2012]. In the hybrid method the state of a group changes deterministically by a partial differential equation (PDE) that describes the dynamics of a continuum of types, while the group-level events (fission and extinction) occur stochastically. The PDE governing population dynamics within the groups is derived from the stochastic rate functions (for births and deaths) by essentially treating the stochastic rates as deterministic rates. The approximation is justified by large population limits, like those in [Champagnat et al., 2006, Puhalskii and Simon, 2012]. In our numerical experiments the groups are fairly large (around 175-200 individuals on average) so the hybrid method is appropriate. Using the PDE (A.1) specified in Appendix 1, the state of a group becomes a continuous density, *z*(*x*), *x* ∈ [0, 1], where 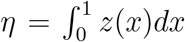 is analogous to *n* (the size of a group) in the Markov chain. The continuous approximation of the within-group population dynamics requires a slight change to (1), which becomes

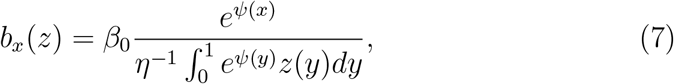

where

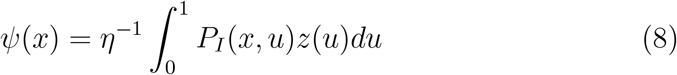

is the expected payoff to an individual with type *x*.

To implement the PDE for a continuum of individual types, we approximate the continuum individual-type space, [0, 1], by a discrete set. We chose a set of 50 equally spaced points between 0 and 1. This leads to a system of 50 nonlinear differential equations, which are derived in Appendix I. The equations are easily and accurately solved by standard numerical techniques.

The group-level events are simulated by discretizing time and determining which (if any) group level events occur in each time interval. Since we are interested in the sample path behavior that leads to an equilibrium (or quasi equilibrium) in our models, and not just the equilibrium itself, this hybrid approach is ideal.

In our model of group-structured populations, every parameter can potentially have direct or indirect effects on every aspect of the evolutionary dynamics. Birth and death rates of individuals generally affect the average group size, but so does the fission rate. Fission and extinction rates affect the number of groups, but so do individual birth and death rates since they affect the group sizes, which affect the fission rates. However, one simplification is that the form of the individual birth rates and the group extinction rates (5), (7) decouples the ecological and evolutionary dynamics in the sense that it ensures the total rates, summed over the individuals or groups, are independent of the games played: the total population-wide extinction rate is always *e*_0_*N* ^2^, and the total group-wide birth rate is always *β*_0_*η*.

### 2.2 Continuous snowdrift and prisoner’s dilemma games

Following [Doebeli et al., 2004], we specify a continuous game by two functions, *C*(*x*) and *B*(*x*), interpreted as cost and benefit functions. In a continuous snowdrift game the payoff function is

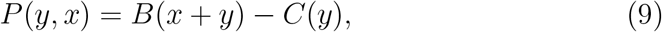

i.e., both players contribute to the benefit *y* receives, but *y* incurs its own cost. A continuous prisoner’s dilemma game has

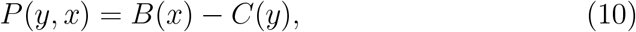

i.e., *y*’s benefit comes only from *x*. In this paper, cost and benefit functions are quadratic, i.e.,

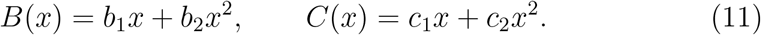

Doebeli et al. [Doebeli et al., 2004] derive conditions, in terms of the quadratic coefficients, for a well-mixed population of individuals playing snowdrift to evolve towards an attracting and evolutionarily stable singular point (an ESS), or to evolve towards an attracting and evolutionarily unstable singular point (a branching point). The details of the algebra are not important here, but we note that both the benefit and the cost function should be increasing on the range of strategies considered (here *x* ∈ [0, 1]), and that *B*^*’*^(0) > *C*^*’*^(0) should hold for *x* to evolve away from 0. It follows for example that if *b*_1_ = 30.0, *b*_2_ = − 7.0, *c*_1_ = 22.8, and *c*_2_ = − 8.0, then the population evolves towards a singular point at *x*^*^ = 0.6, and then branches into two coexisting phenotypic clusters, Figure 1. Similarly, when *b*_1_ = 50.0, *b*_2_ = − 10.0, *c*_1_ = 45.0, and *c*_2_ = − 15.0, the attractive singular branching point is at *x* = 0.50. In the following, we will refer to those parameters as Snowdrift game I and II.

**Figure 1.**
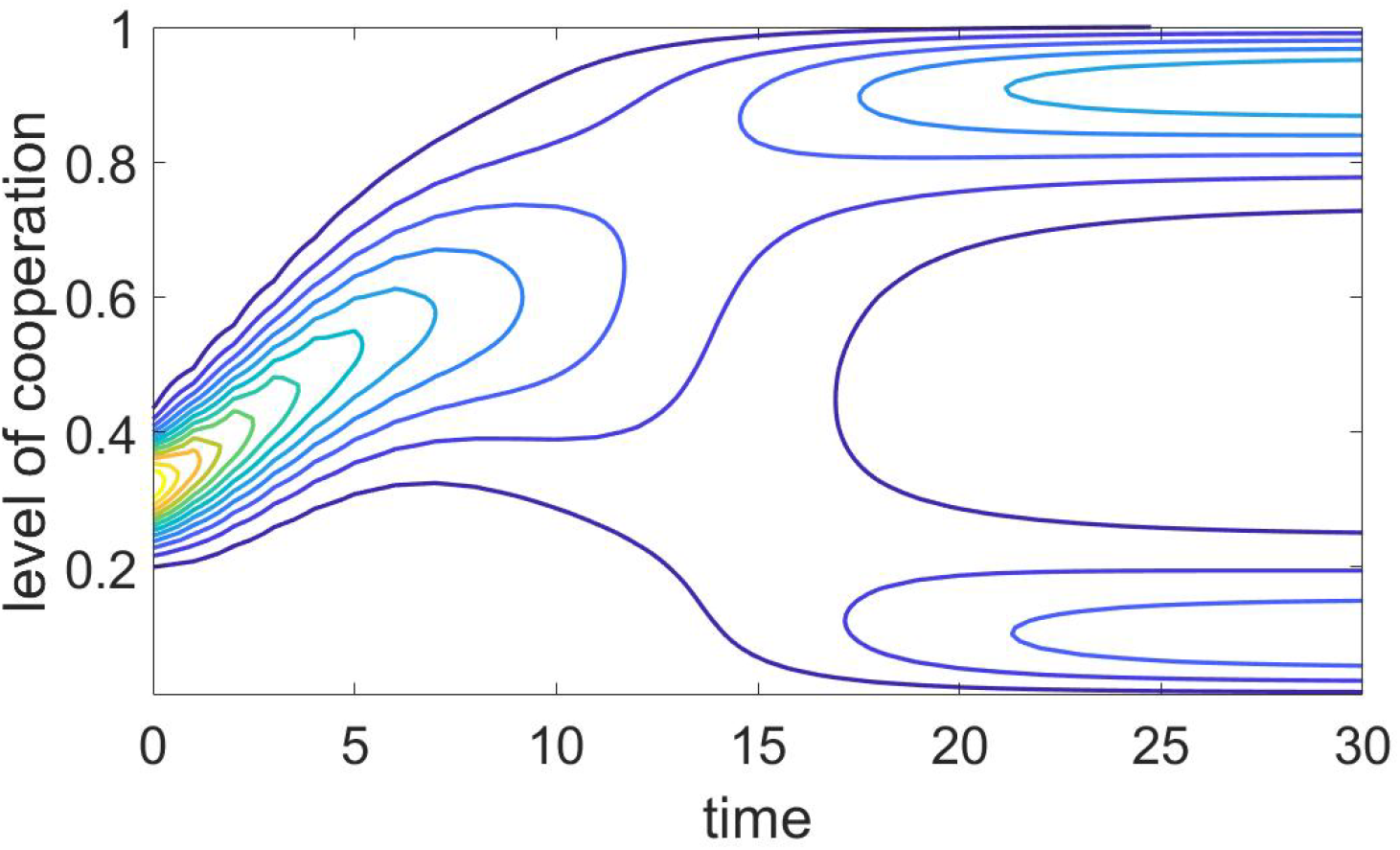
Evolutionary dynamics of individuals playing the Snowdrift game I (Table 1) with evolutionary branching at *x* = 0.6, from (A.1). Note that it takes about 10-20 of our time units to branch. Our simulations are all much longer than that, so there is plenty of time for many generations of groups to branch. However, due to group-level selection pressures, groups that have branched may be less fit, so the population of groups might look very different, e.g., group-level branching may occur.

**Figure 2.**
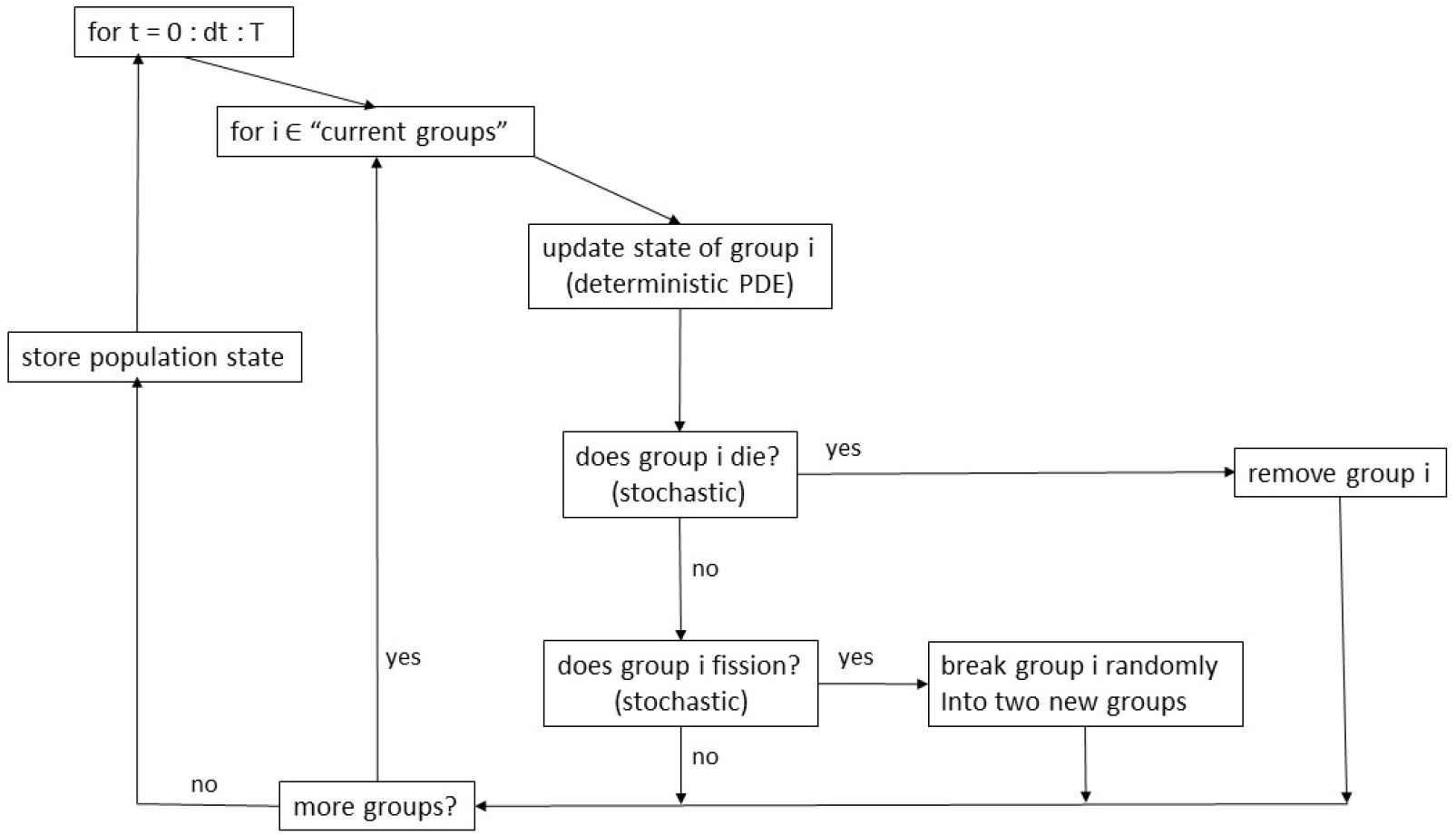
Simulation flow diagram.

The prisoner’s dilemma has much simpler phenomenology, an inevitable evolution towards complete defection, *x* = 0, for any positive *B*(*x*) and *C*(*x*).

**Table 1:** Values of game coefficients used in the simulations.

| Parameter | Snowdrift I | Snowdrift II | Prisoner's |
| --- | --- | --- | --- |
| $b_1$ | 30 | 50 | 1.05 |
| $b_2$ | -7 | -10 | 0 |
| $c_1$ | 22.8 | 45 | 1 |
| $c_2$ | -9 | -15 | 0 |

For simplicity we will use

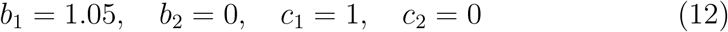

for the prisoner’s dilemma game in our simulations. In Table 1 we summarize the coefficients of the games that are used in this work

The conditions for ESS imply that if the linear and quadratic coefficients of *B*(*x*) and *C*(*x*) are multiplied by the same constant then the resulting evolution is qualitatively (but not quantitatively) the same, e.g., if one branches then the other branches too, at the same singular point. Therefore, without affecting the branching behaviour, we can write the game payoff functions as

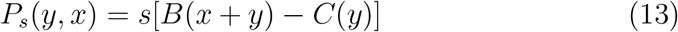

when the game is snowdrift, and

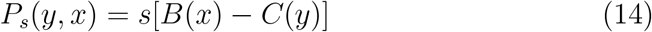

when the game is a prisoner’s dilemma. The parameter *s* ≥ 0 is a scaling factor of the selection pressure.^4^ If *s* = 0 the game is neutral: the players’ strategies have no effect on the outcome of the game, so players all get the same payoff equal to 0. As *s* increases, the payoffs vary more, so there are bigger advantages or disadvantages to different strategies. Thus, increasing *s* makes the game a more important factor in the evolutionary process. We will use the scaling factor in our numerical experiments as follows. To gauge the strength of individual vs group selection, we use identical games at the individual and at the group level, except for the scaling factor *s*, in the examples shown in Figures 3-5. In our last example, the groups play Snowdrift II and individuals play prisoner’s dilemma. At the group level, i.e., for the game between groups, the scaling factor is always set to 1. On the other hand, at the individual level we vary the strength of the given game by varying the scaling factor for any given game. This allows us to determine how the strengths of the games at the individual level (varying *s*) relative to the group level (*s* = 1) affect the evolutionary dynamics at both levels.

**Figure 3.**
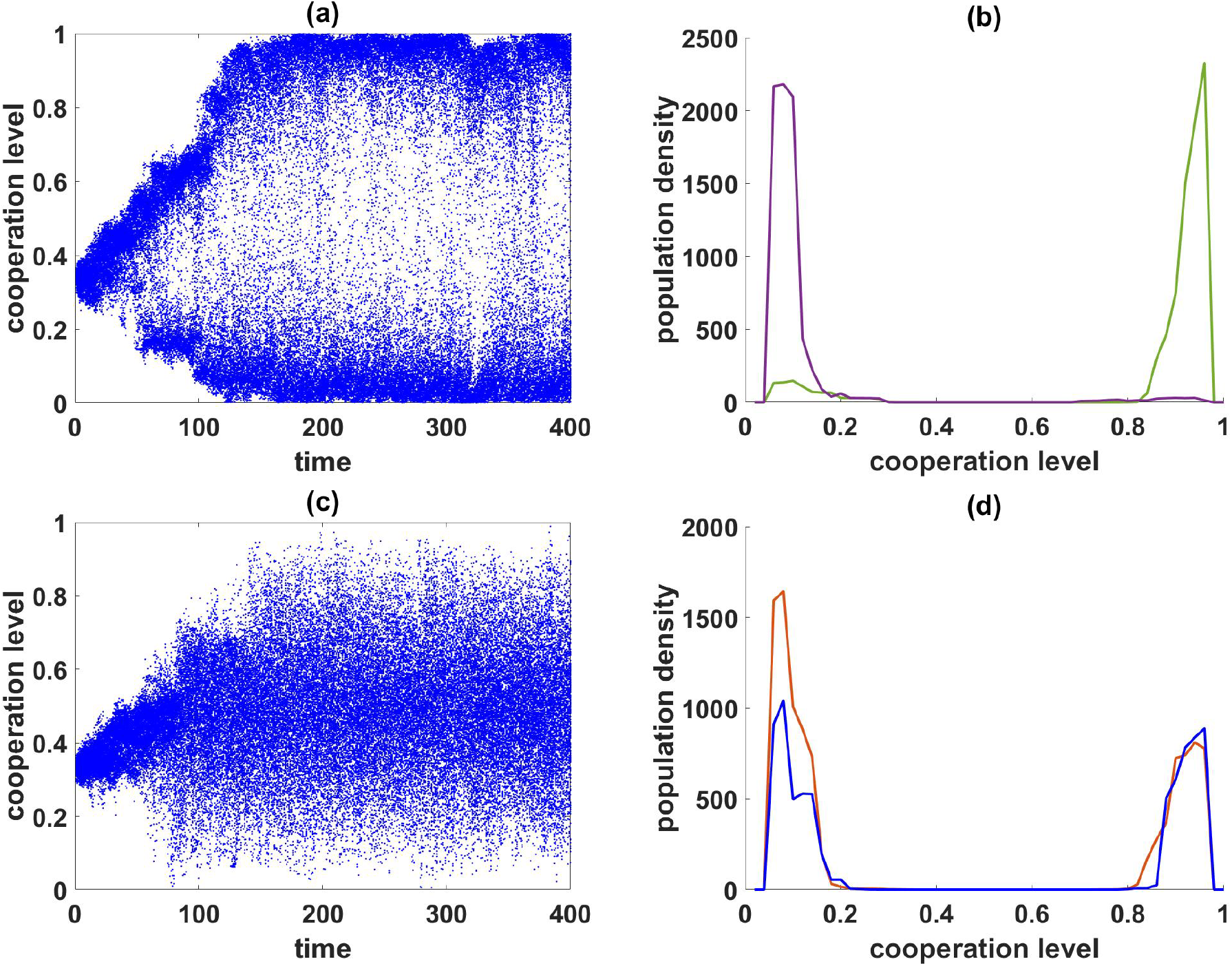
The figure shows dot plots for group-average cooperation levels, *E*(*X*_*t*_ | *G*_*t*_), on the left, and sample group states on the right. The game at both levels is Snowdrift II with the second set of ecological parameters. which branches at *x* = 0.50. Panels (a) and (b) have *s* = .003, where there is clear group-level branching; and panels (c) and (d) have *s* = .010, where there is clear within-group branching (and therefore no group-level branching).

To check robustness of our models and their predictions, we perform simulations for two sets of “ecological parameters”, or rate constants in Eqs. (3-5). That is, for both types of Snowdrift game, I and II, we ran the model with ecological parameters listed in Table 2.

**Table 2:** Values of ecological parameters used in the simulations.

| Parameter | Set I | Set II |
| --- | --- | --- |
| Extinction rate coefficient $e_0$ | $10^{-3}$ | $3 \times 10^{-4}$ |
| Fission rate coefficient $f_0$ | $5 \times 10^{-4}$ | $2 \times 10^{-4}$ |
| Death rate coefficient $d_0$ | $5 \times 10^{-3}$ | $3.5 \times 10^{-3}$ |

### 2.3 Simulations

The simulation works as follows (Figure 2). A time step, *dt*, is chosen small enough so only a small fraction of groups undergo a group-level event in one time step, and also small enough so that the numerical solution of the within-group ODEs is sufficiently accurate. We found that *dt* = 0.10 works well in our examples. Starting with the initial population state at *t* = 0, the simulation updates the population state every *dt* time units up to time *T* (the length of the simulation), which is long enough to see the evolutionary dynamics unfold. In all our experiments we set *T* = 10000, which means there were *T/dt* = 100000 time steps. At each time step, the simulation:

1. updates the state of each group via one iteration of the numerical ODE solver,
2. determines if any group-level events occurred in the time interval, and updates the system state if necessary, and
3. stores important statistics from the new system state for recreating the evolutionary dynamics and constructing plots when the simulation finishes.

The initial state in all our examples is a population of 100 groups, each with 100 individuals with strategies uniformly distributed between 0.25 and 0.35; although the initial state has no effect on the equilibrium.

Simulations of the Markovian model, or the hybrid approximation just described, generate a large amount of data. At each point in time the simulation records the state of the population, Ξ(*t*), which means it records which groups are present, and the current state of each group.

It is not easy to display all this information in the form of simple graphs or tables. For the purpose of observing group-level evolution, including group-level branching, the most informative display seems to be one constructed as follows. Let *X*_*t*_ ∈ [0, 1] denote the type of a randomly chosen individual from the population at time *t*, and let *G*_*t*_ be the identity of the group that individual resides in. Then *E*(*X*_*t*_ | *G*_*t*_) is a random variable whose possible values are all the mean cooperation levels of the groups present at time *t*, i.e.,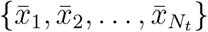. Likewise, *V ar*(*X*_*t*_ | *G*_*t*_) is a random variable whose possible values are the within-group variances. Each group is chosen with probability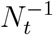, so *V ar*(*E*(*X*_*t*_|*G*_*t*_)) is the variance of the cooperation level of a group picked at random, and *E*(*V ar*(*X*_*t*_|*G*_*t*_)) is the average within-group variance of a group picked at random. Comparing *E*(*V ar*(*X* |*G*)) to *V ar*(*E*(*X* |*G*)) shows the relative strengths of individual-level and group-level selection, e.g., Figure 4. We refer to plots of *E*(*X*_*t*_ |*G*_*t*_), *t* ≥ 0 as “dot plots”, as they show 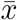 for each group alive at time *t* as a dot. (See Figures 3a, 3c, and 5a-d.) However, the dot plots are not sufficient for describing group states. For that, we plot the population density *z*(*x*), *x* ∈ [0, 1] for representative (randomly chosen) groups, e.g., Figures 3b and 3d.

**Figure 4.**
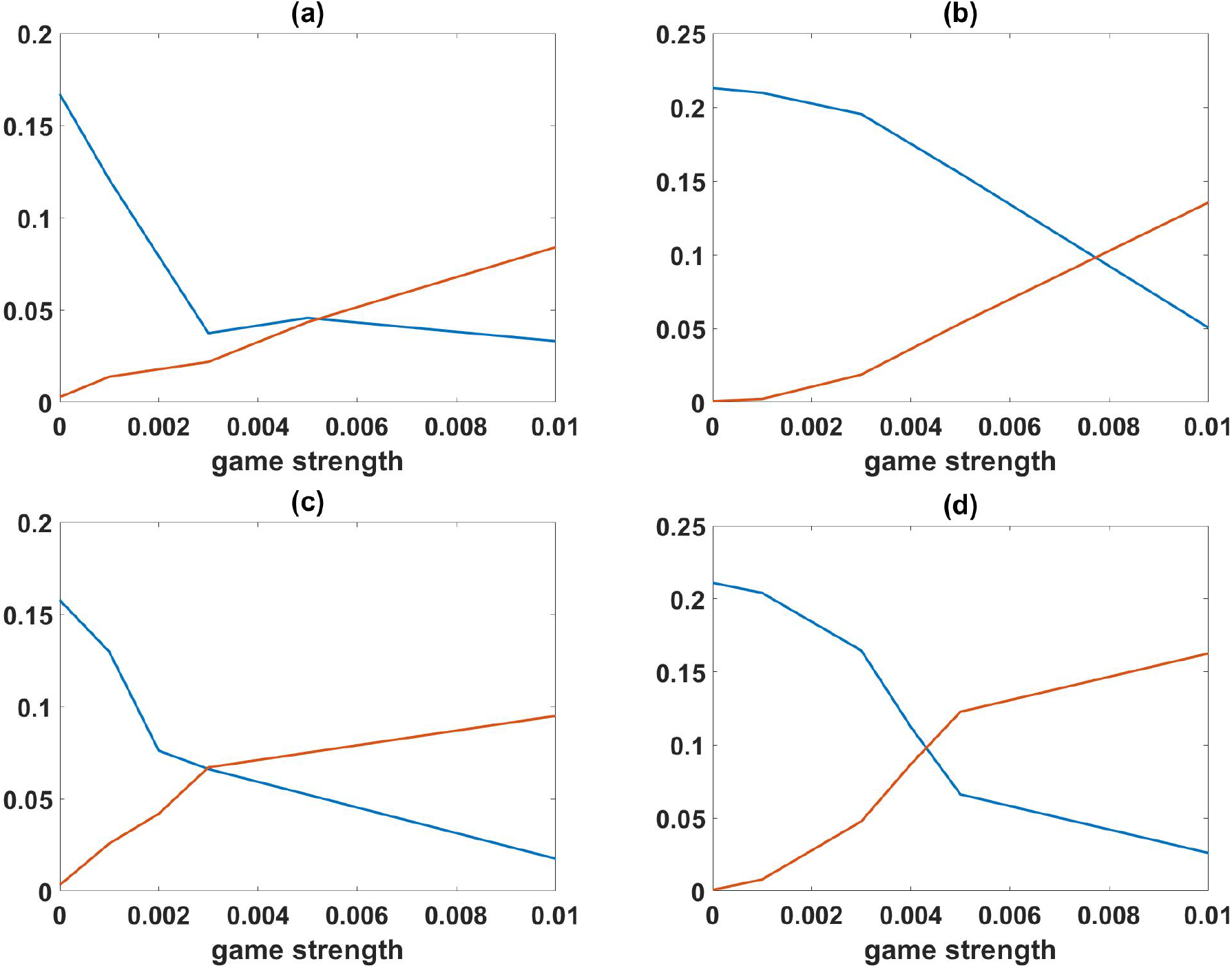
The panels show how the between-group variance *V ar*(*E*(*X* | *G*) (blue) and the within-group variance *E*(*V ar*(*X* | *G*)) (red) depend on the strength of the within-group game, controlled with the parameter, *s* ≥ 0. Panels (a) and (c) have Snowdrift game I and panels (b) and (d) have Snowdrift game II. Panels (a) and (b) are computed using the first set of ecological parameters and panels (c) and (d) are done using the second set.

## 3 Results

There are different ways to show conflict between individual and group level processes in our model, and we have to be content to show only a few representative examples. As a reference point, we chose the group-level game to be the branching Snowdrift game (9, 11) with either the first or second set of coefficients (Table 1). The individual level games vary between Snowdrift games (9,11) and Prisoner’s dilemma (10), as do their relative strengths, *s*.

We begin by showing that group-level branching is possible. When the groups play a branching snowdrift game, while the individuals play a neutral game *s* ≈ 0, the evolutionary dynamics is determined by the selective pressure at the group level, since there is none, or very little, at the individual level. This is shown in Figures 3a,b and 5a,b. In these examples, given enough time every group in isolation would have gradually diversified into a (almost) uniform distribution of types due to random mutational drift (there would be slightly fewer individuals near the boundary in our model because mutations outside of [0, 1] die.) In particular, *E*(*X*_*t*_ | *G*_*t*_) would cluster around 1*/*2. However, the group-level events do not allow this to happen. Instead, *E*(*X*_*t*_ | *G*_*t*_) branches, and the population of groups becomes dimorphic: one branch consist of groups that contain mostly cooperators (high *x*-strategists), and one branch consists of groups that contain mostly of defectors (low *x*-strategists).

We note that adaptive dynamics theory suggests that group-level branching would occur when groups play branching snowdrift, but since some of the basic assumptions of the theory are violated at the group level (in particular, the population of groups is not monomorphic due to fission and within-group dynamics), the demonstration of branching in our model was not assured. It seems to be one of the first examples of evolutionary branching and diversification in continuous traits due to group selection in the literature.

Figure 3c,d shows what happens when the branching snowdrift game is played at both levels, with the game scaling factor *s* > 0 at the individual level. With the within-group game sufficiently strong (e.g., *s* = 0.1), the groups cannot branch, and remain similar in their group-level strategy around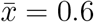. Instead, the branching into cooperators and defectors occurs within the groups.

Thus, as the scaling factor *s* increases and selection exerted by the game at the individual level becomes stronger, the evolutionary dynamics gradually changes from evolutionary branching at the group level, resulting in two different and mostly monomorphic types of groups, to evolutionary branching at the individual level, resulting in all groups being similar and consisting of dimorphic populations of individuals. The transition between the two regimes can be captured by depicting the between group and within group variance at the evolutionary equilibrium as a function of the scaling factor *s*, as is done in Fig. 4. The figure illustrates the crossover in the selective dominance from the the group-level game to the individual-level game as the strength of the latter, *s*, is increased. The crossover happens around the value of *s*^*^ when the stationary between-group variance *V ar*(*E*(*X* | *G*) (blue) becomes less than the average stationary within-group variance *E*(*V ar*(*X* | *G*)) (brown). In the four cases shown in Fig. 4, which represent the combinations between two Snowdrift games and two sets of ecological parameters, the crossover occurs when the strength of individual games exceeds *s*^*^ ≈ 3 × 10^−3^ to 8 × 10^−3^.

In Appendix II we provide a simplistic approximation, based on adaptive dynamics, for the value *s*^*^ of the scaling parameter at which the crossover between dominance of group vs individual selection occurs. While the adaptive dynamics at the individual level is derived as usual, the approximation at the group level is not fully justified, because the group-level evolution involves both individual- and group-level events, and because we neglect the variation generating effects of fissioning. Nevertheless, our approach provides an estimate for the value *s*^*^ at which the rates of evolutionary change in the strategy *x* due to individual-level and group-level selection are approximately the same:

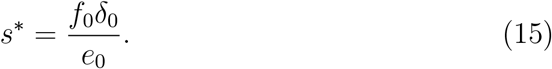

For the parameters used here, this happens at 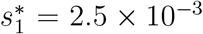 for the first set of ecological parameters and at 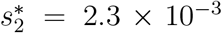 for the second set. While providing useful order of magnitude estimate, these theoretical values are systematically less than the crossover values of *s* observed in Fig. 4. Most likely, the reason for underestimating *s*^*^ is the omission of the effect of randomization of groups during fissioning in the theoretical estimate. As we show in other work [Simon et al., 2024], this effect increases the between group variance, making group-level evolution more effective, which in turn requires stronger individual-level selection, i.e., larger *s*, to balance. We also note that the multi-level partial differential equation models of [Luo, 2014, Cooney, 2019, Cooney, 2020, Cooney and Mori, 2022] could provide an alternative approach to determining the relative strength of the individual-level and the group-level selection regimes, but such models would have to be modified to incorporate games with continuous strategies and nonlinear payoffs.

Figure 5 shows what happens when the between group game remains the branching snowdrift game (9, 11), but the within-group game is a prisoner’s dilemma (10). Branching still occurs when the prisoner’s dilemma game is weak enough, Fig. 5(a,b), but when *s* is sufficiently large, the prisoner’s dilemma forces the population to be less and less cooperative, as one would expect, Fig. 5(c,d).

**Figure 5.**
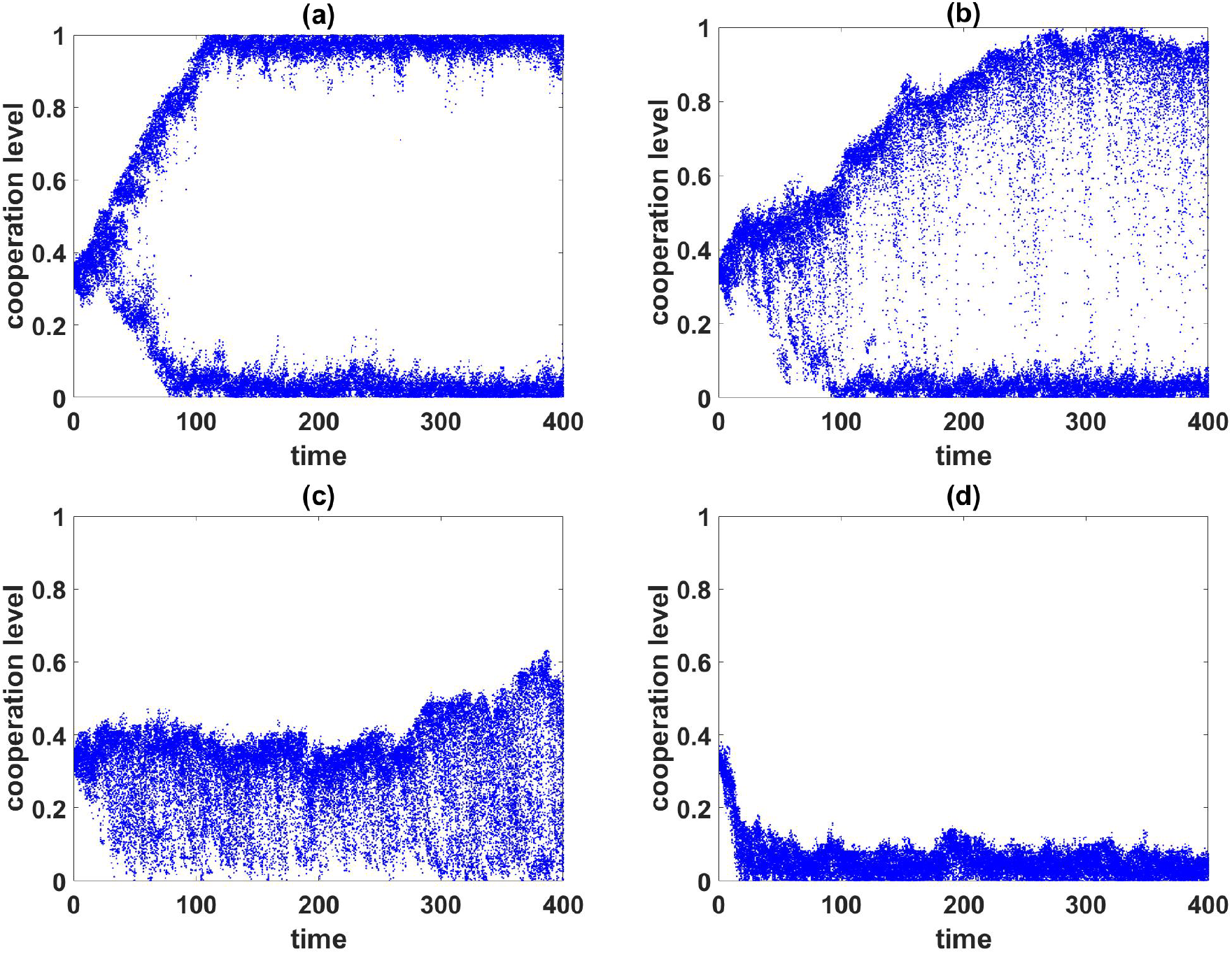
The figure shows dot plots of the group cooperation levels as they change in time for various within-group game strengths in the case where individuals play prisoner’s dilemma, and groups play Snowdrift game II, The first set of ecological parameters is used. In panel (a) the strength of the prisoner’s dilemma is *s* = 0, so group branching occurs unimpeded. As *s* increases, the individual-level selective forces gradually dominate, until cooperators cannot thrive at all. (Panel (b) has *s* = 0.05, panel (c) has *s* = 0.10, and panel (d) has *s* = 0.50.)

## 4 Discussion

Conflict between individual and group level processes allows for the possibility of group selection. Evolution due to group selection occurs when the individuals evolve in a way that is different from how they would have evolved in the same model minus the group-level influences, e.g., fission and extinction [Simon et al., 2013].

Group selection is usually used to explain the evolution of cooperation. In the present paper, we have used the continuous snowdrift game to show that group selection can lead to group-level evolutionary branching, which results in two distinct types of (almost) monomorphic groups – one type of group consisting mostly of cooperators, and the other type of group consisting mostly of defectors – and which occurs despite individual-level selection favouring within-group diversification. We also see group selection facilitating evolutionary branching when individual-level selection favours pure defection.

Usually, group selection is studied using models with “pure” cooperators and “pure” defectors, [Traulsen and Nowak, 2006, Luo, 2014, Simon, 2010, Simon et al., 2013, Simon and Pilosov, 2016]. The model considered here has a richer structure since there is a continuum of cooperation levels instead of just two. Such cases of “continuous public goods games”, in which individuals make variable cooperative investments within groups, and groups use the accumulated individual investments (represented by the average investment in our models) to make cooperative investments in interactions with other groups, can be envisioned to occur in many different biological scenarios. For example, microbes living in microbial mats produce chemicals used by mat mates, and mat structure allows some of these chemicals to diffuse to other mats. Or people pay taxes, which are used for within-group infrastructure, but e.g. also for between group alliances.

Even though evolutionary branching is well-known from the basic continuous snowdrift game, the fact that branching at the group level is possible at all is not obvious. One might think that since groups are “super organisms” they might evolve like ordinary organisms. But the birth and death processes that lead to evolution at the two levels are quite different. In particular, group reproduction occurs when groups fission into pieces. The pieces do not necessarily resemble the parent group at all. Pieces can be considerably smaller than the parent, and can differ significantly in the phenotypic composition.

An interesting, albeit expected, observation was that varying the relative strengths of individual- and group-level selection results in a transition from the between-group to the within-group diversification In the scenario when the effects of individual games are relatively small, the community of groups branches into two classes of groups, one of predominantly cooperators and the other of defectors. At the same time there is no within-group branching: each group consists mostly of cooperators or mostly defectors (even though when left alone, the populations in each group would undergo branching themselves). As the relative selective effect of within-group games is increased, the nature of branching flips: Now the population of each group splits into cooperators and defectors, while the average level of cooperation per group, which defines the outcome of between-group games, becomes similar among groups.

Since all this occurs when the branching snowdrift game is played both at the individual and group level, the non-branching population, whether of individuals in a group or of distinct groups, is in evolutionarily suboptimal state (corresponding to the minimum of invasion fitness). Hence, another way to describe our observation is that evolution at the level at which the selection pressure is weaker does not converge to the scenario that emerges in isolation, yet such an “incomplete” adaptation enables the other level to complete its evolutionary trajectory. Furthermore, we observe that more uniform within-group populations allows groups to better adapt, while diverse within-group population inhibits adaptation at the group level.

Another interesting feature of group selection, which we believe is not limited to this model, is the essentially Lamarckian nature of the evolution of the group phenotype. During the lifetime of a group, the proportions of individual-level phenotypes within the group change over time due to individual-level processes (birth and death). Therefore, the group-level phenotype changes as the group ages. When a group fissions, the phenotypic compositions of the offspring groups depend on the phenotype distribution of the parent group at the time of the fissioning event. How exactly the evolved phenotype distribution in the parent is transmitted to the offspring groups depends on the details of the fissioning process. For example, both offspring distributions could be identical to that of the parent (albeit with smaller numbers of individuals in each offspring group). In our model the fission is essentially random, so in addition to continuous within-group change due to individual-level events, group phenotypes can undergo significant changes due to the fissioning itself. Together, Lamarckian group evolution and fissioning generate the variation at the level of groups upon which group selection can act.

Given the complexity of the two-level selection models considered here, with games with continuous strategies at both levels and the Lamarckian effects mentioned, it is not altogether surprising that the prospects for analyzing our models exactly in the traditional sense (without simulation) is daunting. There are a number of factors that make the analysis difficult. First, the state space, {Ξ}, for the model is infinite dimensional. There is no finite dimensional representation of a group with a continuum of individual types and no bound on the number of individuals it contains, so even the state space for a group is infinite dimensional. There is also no bound on the number of groups in the population. However, the set of possible values of Ξ can be embedded in [0, 1]^∞^, which is a nicely structured mathematical object, so perhaps in that sense the state space is not the primary problem. The real problem is that something analogous to a Markov chain transition matrix for our model (e.g., an infinitesimal generator) is extremely messy. We have described precisely how transitions occur in Section 2, so an infinitesimal generator could be derived in principle. But, while programming simulations of the model is fairly straightforward, constructing the infinites-imal generator appears to be a formidable task. The generator for a model analogous to ours, but with only two individual types (pure cooperators and defectors) was derived in [Puhalskii and Simon, 2012], and the result fills an entire page. We will leave the derivation of an infinitesimal generator, and similar analytical results for the model described here, for future work.

For now, our hybrid simulation/numerical method should be sufficient to study a wide range of multi-level selection scenarios in group-structured populations. It is tempting to generalize the observations from our examples here to explain various biological and social evolutionary scenarios and generate predictions and recommendations. Yet we believe that extensive further studies of different kinds of two-level models and evolutionary setups are necessary to proceed with such generalizations and predictions.

## Acknowledgements

YI acknowledges support from FONDECYT (The National Fund for Scientific and Technological Development of Chile) project no. 1200708, MD acknowledges support from NSERC (The Natural Sciences and Engineering Research Council of Canada) grant no. 219930.

## Appendix

### I. Continuous approximation for individual birth-death process

Here we derive some deterministic approximations for the stochastic birthdeath process within the groups. The approximations are best when the number of individuals in the groups are large enough so that the state of the group at time *t* ≥ 0 is well represented as a continuous density, *z*_*t*_(*x*), *x* ∈ [0, 1]. The change in the number of type *x* individuals between times *t* and *t* + *dt* is due to

1. births from type *x* individuals that do not mutate,
2. births from type *y* individuals that mutate by *x* − *y*, and
3. deaths of type *x* individuals.

We assume that the magnitude of a mutation is determined by a Gaussian with mean 0 and variance *σ*^2^, which results in the following PDE for *z*_*t*_(*x*):

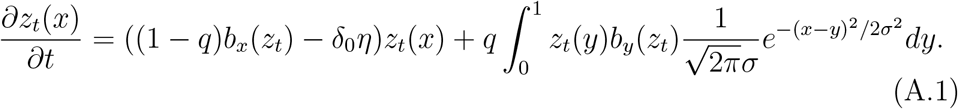

Here

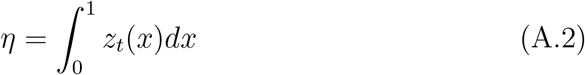

is the number of individuals in the group at time *t*,

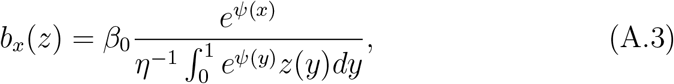

is the birth rate of type *x* in a *z*-group,

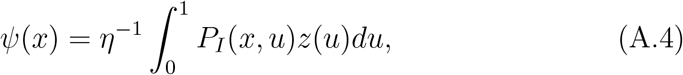

is the expected game payoff, *q* is the probability a birth produces a mutant, *δ*_0_ is the death rate parameter, and *σ* is the mutation size parameter. Note that *b*_*x*_(*z*) and *η* are in fact functionals of the density of individuals, *b*[*z*_*t*_(.), *x*] and *η*[*z*_*t*_(.)], so (A.1) is an integro-differential equation.

This equation can be approximated by a system of nonlinear ODE’s by discretizing [0, 1] into *k* equal pieces. Let *u*_*i*_(*t*) be the number of individuals with type between 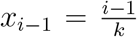 and 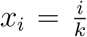, 1 ≤ *i* ≤ *k*. Then (A.1) can be approximated by

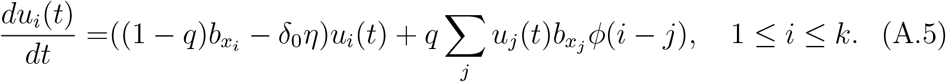

where *φ*(*i* − *j*) is the probability a mutant type *j* is type *i*. This system of ODEs is easily and efficiently solved by standard numerical techniques. In our examples we set *k* = 50. Note that if *σ* is small, as it is in all our examples here, the integral-differential equation (A.1) can be approximated by the diffusion PDE,

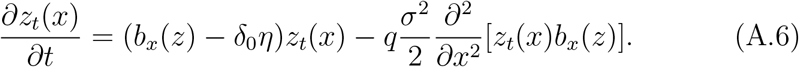

Both (A.1) and (A.6) are consistent with the equations found in [Champagnat et al., 2006], section 4.

### II. Adaptive dynamics estimate for the individual and group evolution rates

Adaptive dynamics is a powerful approximation [Dieckmann and Law, 1996, Champagnat et al., 2002] that is often used to describe the direction and rate of evolutionary change in an evolving population. In particular, adaptive dynamics has been widely used to study the paradigmatic phenomenon of evolutionary branching ([Doebeli, 2011]). The adaptive dynamics analysis of the individual-level snowdrift model presented in [Doebeli et al., 2004]. It shows that the selection gradient *S*_*I*_(*x*) (the gradient of the growth rate of the mutant with type *y* in the presence of a monomorphic resident *x* evaluated at *y* = *x*) for the payoff (13) is

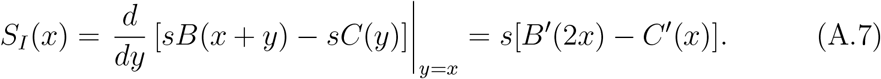

Here we took into account that the birth rate coefficient, *β*_0_, was always set equal to one, *β*_0_ = 1. It follows that the rate of evolution of the strategy *x* within a given group, which has to be assumed monomorphic for the formal applicability of adaptive dynamics, is

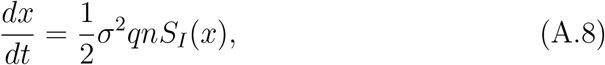

where *σ* is the variance of the mutational distribution, *q* is the probability of mutation at birth, and *n* is the size of the group (see [Dieckmann and Law, 1996] for details).

Since the group-level evolution depends on events occurring at both the individual and the group level, the adaptive dynamics of group strategies is less clear. Nevertheless, we can derive a simplified version of the group adaptive dynamics by assuming that selection is neutral at the individual level (*s* = 0 in the individual game, Eqs. (9,10)). We first note that that the invasion fitness 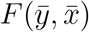 for a few “mutant” groups of type 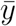 into a resident population of 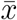 groups is simply the difference between the fission rate and the death rate of 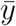 groups in the resident population:

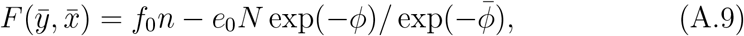

where 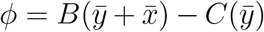 is the payoff of a mutant group 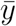 playing with a resident one, 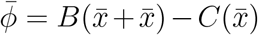 the payoff of a resident group playing with another resident group (identical to it), *N* is number of resident groups, and *n* is the size of the mutant 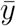 group. Because the group fissioning events are rare on the scale of individual (within-group) population dynamics, *f*_0_ ≪ *δ*_0_, *n* is assumed to be the same for all groups. Setting 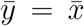 in the above expression and taking into account the definition of the group allows us to calculate

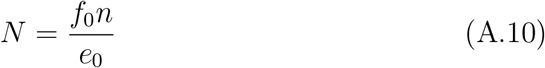

(because the growth rate of the resident must be 0). Moreover, at the within-group equilibrium (which for *f*_0_ ≪ *δ*_0_ establishes itself quickly after each fissioning event) we have *n* = 1*/δ*_0_. The formally computed group-level selection gradient is then

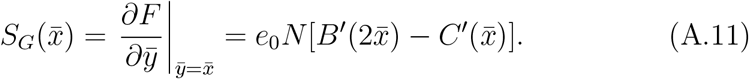

Here we took into account that groups play games with *s* = 1. Similarly to (A.8), we attempt to write the adaptive dynamics equation for the group strategies as

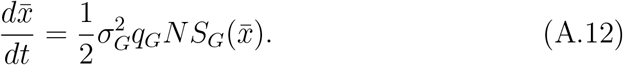

However, the meaning of the constants 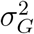, the variance of 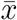 during fis-sion, and *q*_*G*_, the probability of modifying 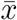 during fission, is very different from that of the corresponding terms in the adaptive dynamics of the individual-level process. First, the changes in 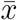 accumulate continuously during a group’s lifetime due to mutations in the individuals comprising the group. This reflects the Lamarkian nature of the evolution of group phenotypes that we have discussed in the main text. While the derivation of adaptive dynamics assumes instantaneous mutations at birth, we can still use, as an approximation, the adaptive dynamics philosophy replacing the instantaneous mutations at birth by the mutations accumulated between consecutive fission events. This period Δ*t* is on average equal to the inverse fission rate, so from (4)

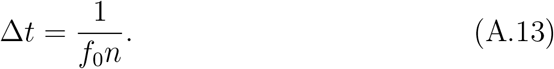

During Δ*t* an average individual accumulates mutational effects with variance

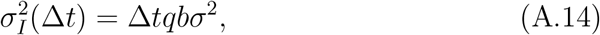

where *b* = 1 is the average individual birth rate. Taking into account that the group type is the average of individual types in the group,

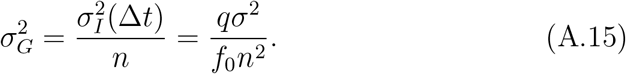

Because individual mutations are occurring continuously within groups over the time span Δ*t*, the probability of mutation of the group phenotype is one, *q*_*G*_ = 1.

Overall, the rates of evolution due to group selection is different from that due to individual selection by the scaling factor *χ*

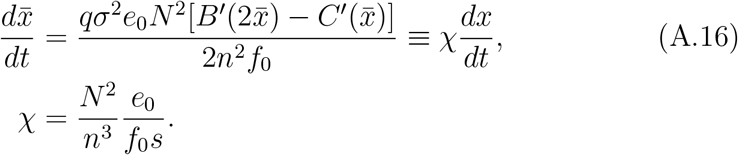

The expression for *χ* can be further simplified using *N* = *f*_0_*n/e*_0_ and *n* = 1*/δ*_0_. Qualitatively, the transition from the individual-level to the group-level selection regime should happen for *s*^*^ for which *χ* = 1,

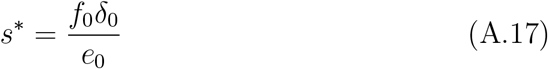

For example, for the first set of ecological parameters in Fig. 4(a,c) *e*_0_ = 0.001, *f*_0_ = 0.0005, *δ*_0_ = 0.005 this happens for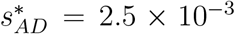, which is less than the crossover point observed in the simulations, where the crossover happens around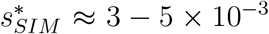. Hence, while the two-level adaptive dynamics considered here does provide a qualitatively reasonable approximation for salient features of the two-level evolutionary process, there exist other factors that are not accounted for in the simple estimate. The most plausible factor responsible for the underestimation by the adaptive dynamics of the crossover value, 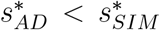 is that random fissioning of groups produce an additional contribution to between group variation, which enhances the effectiveness of group-level evolution and requires a stronger individual-level selection (larger *s*) to balance it. We do not consider this effect here, and instead present a more detailed analysis of various types of fissioning in another article [Simon et al., 2024].

## Footnotes

1 Department of Mathematical and Statistical Sciences, University of Colorado Denver

2 Department of Physics, Center for Interdisciplinary Research in Astrophysics and Space Science, University of Santiago of Chile

3 Department of Zoology and Department of Mathematics, University of British Columbia

4 This is not a weak selection analysis as *s* does not have to be small.

